# The Effective Visual Acuity of Butterflies in Flight

**DOI:** 10.64898/2026.09.30.755797

**Authors:** Jack A. Supple, Wenxin Li, Marko Ilić, Monika Kos, Huai-Ti Lin, Gregor Belušič, Holger G. Krapp

## Abstract

The spatial visual acuity of an eye is determined by both the spatial sampling density of the retina and the angular sensitivity of individual photoreceptors. In insect compound eyes, these are respectively characterised by the interommatidial angle, Δφ, and photoreceptor acceptance angle, Δρ. During movement, however, finite photoreceptor integration times cause retinal image motion to broaden the effective acceptance angle, potentially degrading spatial resolution beyond its anatomical limit. This problem is especially acute in flying insects, which experience large rotational velocities during locomotion and must actively stabilise gaze to limit retinal image motion. Butterflies present a particularly demanding case because their flight is characterised by large, wingbeat-coupled thoracic oscillations. Here, we quantify how gaze stabilisation limits motion blur in freely flying Monarch butterflies (*Danaus plexippus*) by combining free-flight kinematic reconstruction of the head and thorax with separate intracellular measurements of photoreceptor acceptance angles and integration times from quiescent butterflies. We found that, without head compensation, thoracic oscillations would broaden the effective photoreceptor acceptance angle from its quiescent value of 2.17° to 9.38° during take-off. Compensatory head movements substantially reduced this motion-induced broadening, limiting the effective acceptance angle to 5.28°. Nevertheless, this 2.4-fold broadening of effective photoreceptor acceptance angle represents a substantial degradation of spatial acuity in flight, highlighting motion blur as an important constraint on the visual ecology of flying butterflies.

## Introduction

Biological vision is costly, both in material investments^1^ and energetic expenditure^2–4^. In blowflies, for example, photoreceptors operating under daylight conditions consume energy equivalent to approximately 8% of resting metabolic consumption^2^, while photoreceptors capable of transmitting higher information rates incur disproportionately greater energetic costs^5^. Such costs impose strong selective pressure for efficient allocation of visual resources. Visual investment is therefore typically regionalised across the eye, with local variation in spatial acuity, temporal resolution, luminance sensitivity, and spectral tuning depending on species-specific sensory and behavioural ecology.

Spatial visual acuity is jointly determined by the sampling density of the eye and the degree of spatial blurring by individual photoreceptors^6^. In insects, these are characterised respectively by the interommatidial angle, Δφ, and the photoreceptor angular acceptance width, Δρ^7,8^. Finer spatial resolution requires smaller values of both, and visual performance is maximised when the optical resolution of individual photoreceptors is Nyquist-matched to the sampling density of the ommatidial array^7^. For the hexagonal compound eye lattice, this matching occurs when Δρ is 1.7–2 times Δφ (depending on hexagonal sampling geometry)^7^. Light-adapted diurnal insect eyes, however, are typically under-sampled with Δρ≈Δφ^8^, such that the spatial resolution of the ommatidial array is insufficient to sample the smallest detail resolvable by an individual photoreceptor^8^.

However, these anatomical limits describe spatial resolution only for a stationary retinal image^7^. During movement, image motion across the retina introduces an additional source of spatial blur due to the finite photoreceptor temporal integration time, Δt^9^. During this integration period, a point source moving at angular velocity ω traverses an angle ωΔt, effectively broadening photoreceptor angular acceptance beyond its anatomically defined width, Δρ^7^. Consequently, the effective spatial resolution of the eye depends not only on the anatomical distributions of Δρ and Δφ, but also on photoreceptor temporal dynamics and retinal image velocity. This motion blur has been proposed to account for the apparent under-sampling of the insect compound eye, bringing the effective Δρ/Δφ sampling ratio closer to optimal^8,10^.

Crucially, however, retinal image velocity can be actively reduced by stabilising the orientation of the visual system during self-motion^11^, thereby limiting motion blur and preserving spatial resolution closer to its anatomical limit^7^. In flying insects, compensatory movements of the head provide the primary mechanism for reducing retinal image motion. Gaze stabilisation has been extensively characterised in insects, with tethered experiments quantifying dynamic range^12–15^ and multisensory inputs^14,16–18^, while free-flight measurements detail how coordinated head movements actively structure retinal image motion through rapid gaze shifts interspersed with periods of relative stability^19–26^. However, the extent to which gaze stabilisation reduces motion blur, and thereby sets the spatial information available in flight, remains less well described.

Butterflies present an especially demanding case for gaze stabilisation owing to their unusually erratic style of flight^27,28^. Their characteristically large wings relative to body size favour low-frequency (∼10-20 Hz^29^), high-amplitude wing strokes that generate substantial oscillatory forces and torques on the body in flight^30–32^. Pitching body oscillations help redirect aerodynamic forces toward downstroke weight support and upstroke forward propulsion, improving flight power efficiency^33–35^. However, these same oscillations generate large thoracic angular velocities that, in the absence of compensatory head movements, would produce substantial retinal image motion and motion blur.

Here, we quantify the kinematic demands and visual consequences of gaze stabilisation in freely flying monarch butterflies (*Danaus plexippus*). We first characterise 6-degree-of-freedom (6-DOF) thorax kinematics during take-off and cruising flight, before reconstructing head movements during take-off to determine how effectively these rotations are attenuated at the retina. Finally, we analyse the resulting head kinematics to estimate retinal image motion and its impact on effective spatial acuity during flight.

## Results

### Thoracic pitch oscillations dominate take-off and cruising flight kinematics

We first characterised the thorax kinematics of flying Monarch butterflies to determine the stabilisation demands placed on the neck motor system. We compared thorax motion during take-off from a perch with that during forward cruising flight in an indoor arena (Figure 1A). Example trajectories indicate that both behaviours involve substantial changes in thorax orientation during translational motion (Fig. 1B–E). Take-off trajectories were more vertically directed and had lower total speeds (median = 0.68 m·s⁻¹, inter-quartile range, IQR = 0.51-0.86 m·s⁻¹, n = 43 trajectories, 11 animals) than cruising flight trajectories (median = 1.08 m·s⁻¹, IQR = 0.87-1.35 m·s⁻¹, n = 1194 trajectories from 10 animals; Fig. 1Fi). Take-off was correspondingly associated with lower horizontal velocity magnitudes in global coordinates (take-off median = 0.49 m·s⁻¹, IQR = 0.30-0.67 m·s⁻¹; cruising median = 0.88 m·s⁻¹, IQR = 0.60-1.19 m·s⁻¹; Fig. 1Fii), while vertical velocity magnitudes were similar between the two flight behaviours (take-off median = 0.37 m·s⁻¹, IQR = 0.13-0.59 m·s⁻¹; cruising median = 0.52 m·s⁻¹, IQR = 0.29-0.73 m·s⁻¹; Fig. 1Fiii). Projection of the total velocity vector into the butterfly’s thoracic frame of reference showed that cruising flight was dominated by motion along the thoracic frontal (x) axis (Fig. 1Gi-iii), whereas take-off flight showed substantial components along both the thoracic frontal (x) and dorsal (z) axes (Fig. 1Gi-iii), consistent downstroke lift production^36^.

**Figure 1:**
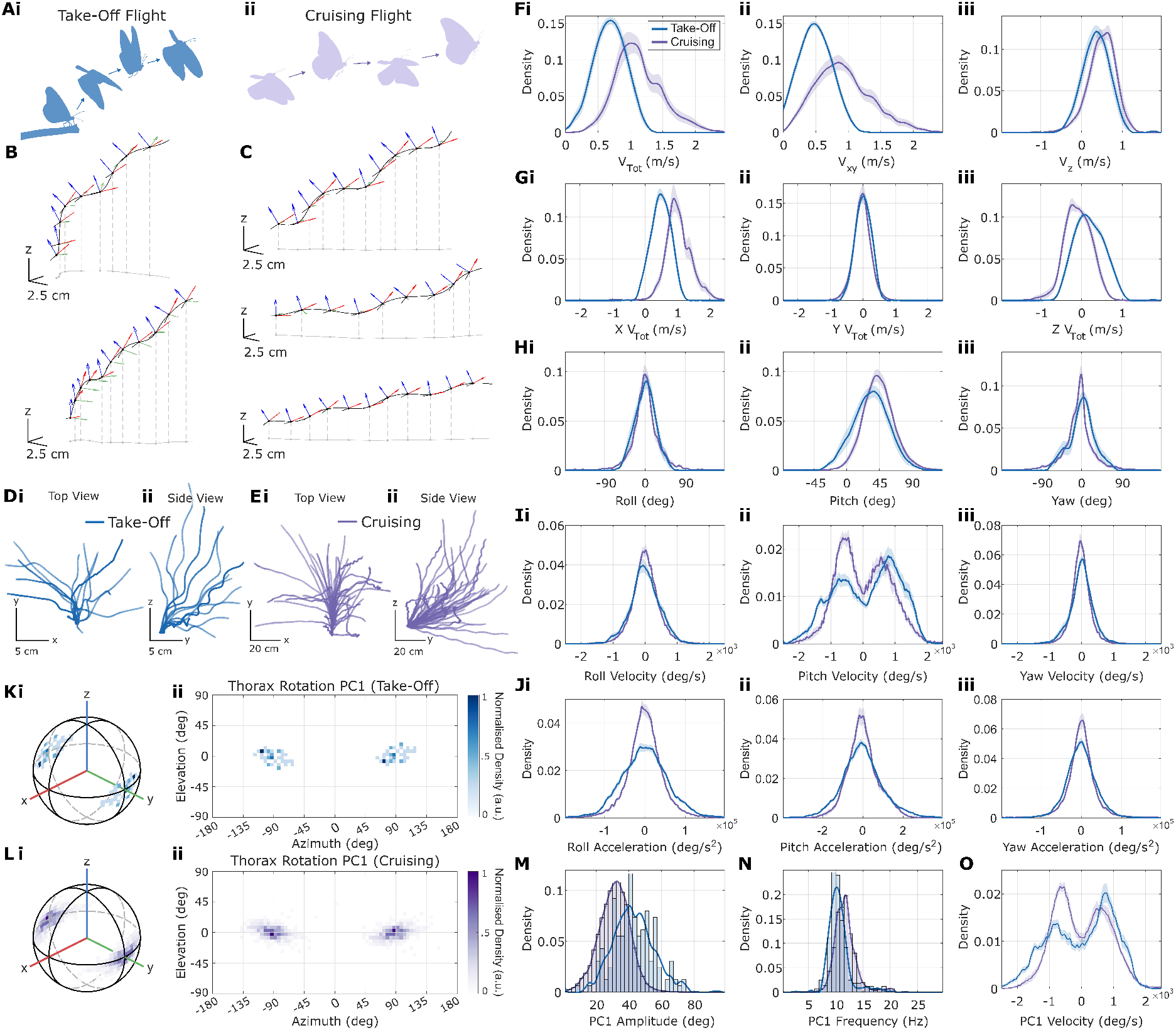
Monarch Butterfly Thorax Kinematics during Take-Off and Forward Cruising Flight. **(A)** Butterfly thorax kinematics during (i) take-off from a perch, and (ii) cruising flight within an indoor flight arena. **(B-C)** Example thorax trajectories for (B) take-off, and (C) cruising flight. Continuous black line shows thorax centre-of-mass. Red/green/blue vectors represent meso-metathoracic frontal (x), lateral left (y), and dorsal (z) axes, respectively. Axes are displayed at 28 ms intervals. Black dots and tails represent the butterfly thorax axis. Grey lines show projection of the 3D trajectory onto the global xy plane (but for clarity do not necessarily indicate ground location). **(D-E)** Top (i) and side (ii) views of example trajectories for (D) take-off and (E) cruising flight. Trajectories are aligned by their initial horizontal velocity (in global coordinates). Only 50/1194 cruising trajectories are shown for clarity. **(F)** Thorax translational velocity histograms for take-off (blue) and cruising flight (purple). (i) Total velocity magnitude, V_tot_; (ii) Horizontal velocity component magnitude, V_xy_; (iii) Vertical velocity component magnitude, V_z_. **(G)** Total translational velocity projected onto each thoracic axis. (i) Projection onto thoracic frontal (x) axis; (ii) lateral left (y) axis; and (iii) dorsal (z) axis. **(H)** Thorax orientation histograms relative to global coordinates: (i) roll; (ii) pitch; (iii) yaw. **(I)** Histograms of rotational velocity around each thoracic axis: (i) roll; (ii) pitch; (iii) yaw. **(J)** Histograms of rotational acceleration around each thoracic axis: (i) roll; (ii) pitch; (iii) yaw. **(K-L)** First principal component (PC1) of thorax rotation for (K) take-off, and (L) cruising flight. PC1 is calculated separately for each flight trajectory, and both poles of the PC1 rotation axis are presented as histograms in (i) spherical coordinates, and (ii) cylindrical projections. Colormap represents normalised histogram density. **(M)** Peak-to-peak amplitude of thorax rotation about PC1. **(N)** Peak-to-peak frequency (oscillations per second, Hz) of thorax rotation about PC1. **(O)** Histograms of thorax rotational velocity about PC1. For all histograms (F-M): Take-off N=43 trajectories from 11 animals, cruising flight N=1194 trajectories from 10 animals.

Relative to the global coordinate frame, thorax roll orientation was symmetrically distributed about 0**°** with IQRs of −16.54-16.05**°** and −14.07-16.05**°** for take-off and cruising flight, respectively (Figure 1Hi). In contrast, thorax pitch had a median of 34.0**°** during take-off (IQR: 16.87-50.67**°**), and was slightly more elevated during cruising flight with a median of 43.3° (IQR: 29.60-58.02°; Figure 1Hii). Yaw orientation, relative to initial heading, was slightly more variable for take-off flights compared with cruising flights, with an IQR of −14.57-20.49° and −20.98-8.64**°**, respectively (Figure 1Hiii).

Roll and yaw rotational velocity distributions were similar in take-off and cruising flight, with both centred at approximately 0 °·s⁻¹ (Figure 1Ii,iii). Roll velocity IQR was −278.21-290.22 °·s⁻¹ during take-off and −230.17-250.19 °·s⁻¹ during cruising flight (Figure 1Ii), while yaw velocity IQR was −190.14-218.16 °·s⁻¹ and −182.14-146.11 °·s⁻¹ (Figure 1Iiii), respectively. In contrast, both take-off and cruising flight pitch velocity distributions were bimodal, reflecting the thoracic pitching oscillation experienced with every wing-stroke (Figure 1Iii). Pitch velocity distributions peaked at approximately −630 °·s⁻¹ and +830 °·s⁻¹ for take-off flights, and −540 °·s⁻¹ and +570 °·s⁻¹ for cruising flights (Figure 1Iii). Pitch velocities were slightly broader for take-off flight, with IQR of −682.51-914.68 °·s⁻¹, compared with −622.46-642.48 °·s⁻¹ for cruising flight. Rotational acceleration distributions were also broader in take-off than in cruising flight, with roll acceleration IQRs of −3.82-3.58 x10^4^ °·s⁻^2^ (take-off) and −2.42-2.42 x10^4^ °·s⁻^2^ (cruising), pitch acceleration IQRs of −6.61-5.57 x10^4^ °·s⁻^2^ and −4.76-4.28 x10^4^ °·s⁻^2^, and yaw acceleration IQRs of −2.30-2.14 x10^4^ °·s⁻^2^ and −1.74-1.74 x10^4^ °·s⁻^2^, respectively (Figure 1J).

These initial analyses suggested that thoracic pitching oscillations are the dominant kinematic component during both take-off and cruising flight. To quantify this more directly, we next extracted the principal rotational component of each trajectory using singular value decomposition (Figure 1K-L). For both take-off and cruising flight, the first principal axis of rotation (PC1) was closely aligned with the thoracic pitch axis (Figure 1K-L). The peak-to-peak amplitude of rotation about PC1 was greater during take-off than during cruising flight (take-off median = 42.30°, IQR = 34.48-50.61°; cruising median = 31.55°, IQR = 25.20-37.66°; Figure 1M). Oscillation frequency (corresponding to the wing-stroke frequency) was slightly lower during take-off (median = 10.30 Hz, IQR = 9.40-11.36 Hz) compared with cruising flight (median = 11.58 Hz, IQR = 10.45-12.63 Hz; Figure 1N). The PC1 rotational velocity distribution closely resembled the bimodal pure-pitch velocity distribution, which are broader during take-off (take-off maxima: −802.60 °·s⁻¹ and +738.60 °·s⁻¹, IQR = −810.60-818.61 °·s⁻¹; Figure 1O) compared to cruising flight (cruising maxima: −606.50 °·s⁻¹ and +642.48 °·s⁻¹, IQR = −666.49-658.49 °·s⁻¹; Figure 1O).

### Gaze stabilisation during take-off flight

The extent to which these large thoracic rotations translate into retinal image motion depends on how effectively they are compensated by movements of the head. We therefore reconstructed head orientation during take-off flight using the stereotyped natural white spot markings on the head capsule of *D. plexippus*, which provided sufficient contrast for stereo-videographic tracking (Figure 2A; Figure S1). Monarch butterflies maintained a stable head orientation throughout take-off, despite the large amplitude thoracic oscillations introduced with each wing-stroke (Figure 2B-D). Relative to the thorax, the head was pitched on average with a slight ventral tilt of −14.24° (median; IQR: −26.14 to −3.06°; Figure 2E-F), corresponding to a median orientation in global coordinates of 15.89° (IQR = 4.13-28.62°; Figure 2G). Average head roll and yaw orientation relative to the thorax was approximately symmetric around 0° and less variable than pitch (roll median = −0.36°, IQR = −9.01-8.29°; yaw median = 1.80°, IQR = −5.41-9.01°; Figure 2F).

**Figure 2:**
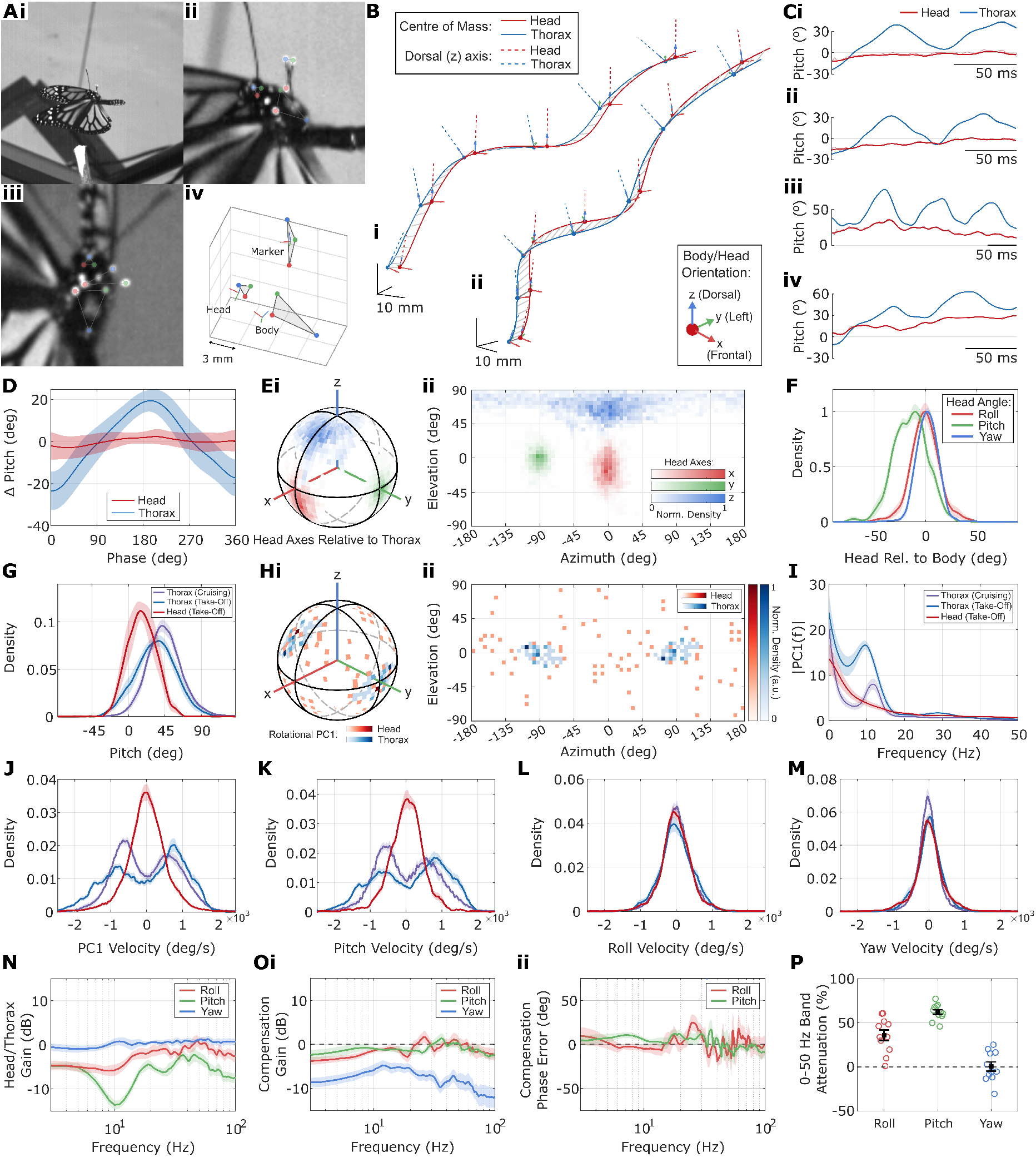
Monarch Butterfly Head Stabilisation during Take-Off Flight. **(A)** Head kinematics were reconstructed during take-off flight by tracking the natural white spot patterning on the dorsal head capsule. (i) Example cropped view of a Monarch butterfly taking off from a perch. Note the carbon-fibre frame affixed to the meso-metathorax, which is used to estimate body axis. (ii-iii) Example cropped stereo image pair, with colored highlights indicating points used to track the head and carbon-fibre frame. Also indicated is the triangle used to define the thorax axis, formed by the tegulae and the dorsal tip of the scutellum. Note that since the tegulae move with each wing-stroke, this reference triangle was measured only at the maximum downstroke depression for all animals. (iv) 3D reconstructed reference points, and coordinate axes of the head, carbon-fibre marker, and thorax. Red/green/blue vectors represent frontal (x), lateral left (y), and dorsal (z) axes, respectively. **(B)** Example trajectories showing the centre of mass and coordinate axes of the head (red) and thorax (blue). Red/green/blue vectors represent frontal (x), lateral left (y), and dorsal (z) axes, respectively. **(C)** Example time course of body and head pitch angles relative to the global horizontal plane. **(D)** Wing stroke-averaged normalised pitch rotation (Δ pitch) of the body (blue) and head (red). Oscillation phase is defined relative to thoracic pitch minima. Both head and body pitch angles are mean-subtracted to show angular changes, rather than their absolute values. n = 48 wing strokes from 7 animals. Mean ± Std. **(E)** Head axis distributions relative to the thorax. Red/green/blue represent density histograms of relative head frontal (x), lateral left (y), and dorsal (z) axes, respectively. Normalised density histograms are presented in (i) spherical coordinates, and (ii) cylindrical coordinates. **(F)** Head axis distributions relative to the thorax. Head roll (red), pitch (green), and yaw (blue) orientations are colored according to the axis of rotation illustrated in (E). Note that here roll (red), corresponds to rotation about the x axis colored red in E. **(G)** Global pitch histograms during take-off for the head (red), and thorax (blue), and during cruising flight (purple). **(H)** First rotational principal component (PC1) distributions of the head (red) and thorax (blue). Thorax PC1 distribution reproduced from Figure 1. (i) Spherical coordinates, (ii) cylindrical coordinates. **(I)** Fourier transform of rotation about PC1 during take-off for the head (red), and thorax (blue), and during cruising flight (purple, note cruising PC1 defined in Figure 1). **(J)** Rotational velocity histograms about PC1 during take-off for the head (red), and thorax (blue), and during cruising flight (purple). **(K-M)** Rotational velocity histograms about (K) pitch, (L) roll, and (M) yaw during take-off for the head (red), and thorax (blue), and during cruising flight (purple). **(N)** Bode plot showing thorax-to-head transfer functions for head roll (red), pitch (green) and yaw (blue) orientation during take-off flight. Negative gain indicates attenuation of thoracic rotation at the head. **(O)** Bode plot showing (i) compensation gain and (ii) phase error for roll (red), pitch (green) and yaw (blue) during take-off flight. A compensation gain of 0 dB indicates equal amplitude compensatory movements. A phase error of 0° indicates anti-phase compensatory movements. Note that phase error for yaw is highly variable and omitted for clarity. **(P)** Percentage attenuation of thoracic movements by the neck motor system for roll (red), pitch (green) and yaw (blue) during take-off flight

In contrast to the pitch-dominated thoracic oscillations, the principal axis (PC1) of head rotation was more broadly distributed in azimuth (Figure 2H). Fourier analysis of head PC1 motion showed that the prominent 10–12 Hz oscillations observed in the thorax were strongly attenuated at the head (Figure 2I). Consistent with this, the head PC1 rotational velocity distribution was unimodal with a median of 2.00 °·s⁻¹ and an IQR of −306.23-314.23 °·s⁻¹, rather than featuring the broader bimodal distribution observed for the thorax (Figure 2J). Whilst the distribution of head pitch velocities was similarly attenuated relative to the thorax (pitch velocity median = 30.02 °·s⁻¹; IQR = −262.19-306.23 °·s⁻¹; Figure 2K), head roll and yaw velocity distributions followed those of the thorax more closely (roll velocity median = −10.01 °·s⁻¹; IQR = −246.18-250.19 °·s⁻¹; yaw velocity median = 2.00 °·s⁻¹; IQR = −202.15-214.16 °·s⁻¹; Figure 2L-M).

We quantified thorax-to-head transfer functions to determine the extent to which thoracic rotation appeared in global head orientation (Figure 2N). Pitch rotations were attenuated across most frequencies, with a pronounced minimum in head/thorax gain at the dominant ∼10 Hz thoracic pitch oscillation (Figure 2N). Roll attenuation was present but less pronounced, while yaw gain remained closer to 0 dB (Figure 2N). Correspondingly, pitch and roll compensation gains and phase errors approached 0 dB and 0°, respectively (Figure 2O), indicating that approximately equal-amplitude, anti-phase head rotations counteracted thoracic oscillations. Yaw compensation gain was substantially lower and phase estimates more variable (Figure 2O), consistent with the small amplitude of compensatory yaw rotations. Across 0–50 Hz, thoracic rotation amplitudes were attenuated at the head by 36.0 ± 6.0% in roll, 62.3 ± 2.7% in pitch, and 0.5 ± 5.2% in yaw (mean ± SEM, N = 11 animals; Figure 2P).

### Motion blur substantially degrades effective visual acuity in flight

Our kinematics data indicate that the butterfly gaze stabilisation system suppresses a substantial portion of the thoracic pitching oscillations experienced during flight. Nonetheless, the rotational velocity distribution of the head remains substantial, with a median absolute pitch velocity of 300.75 °·s⁻¹ and 95^th^ percentile of 997.56 °·s⁻¹. These rotational velocities indicate that substantial retinal image motion persists despite head stabilisation, with potential to degrade spatial visual acuity.

We assessed this by considering how head velocity degrades the effective visual acuity of the eye via motion blur. The intrinsic spatial resolution of the eye is determined by two key parameters (Figure 3A): the inter-ommatidial angle, Δφ, and the photoreceptor angular acceptance, characterised by width Δρ, which determines the degree of spatial blurring prior to sampling^7^. Visual motion degrades spatial resolution because photoreceptors integrate light over a finite time window, characterised by an integration time, Δt^7,9^. During this integration time, a point moving at angular velocity, *ω*, sweeps through an angular distance *ω*Δ*t*, thereby introducing additional motion-dependent blur^7^. Approximating the angular acceptance function and motion-blur function as Gaussians^7^, their convolution produces an effective angular acceptance function with width, *Δρ_eff_*, given by:

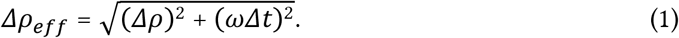

**Figure 3:**
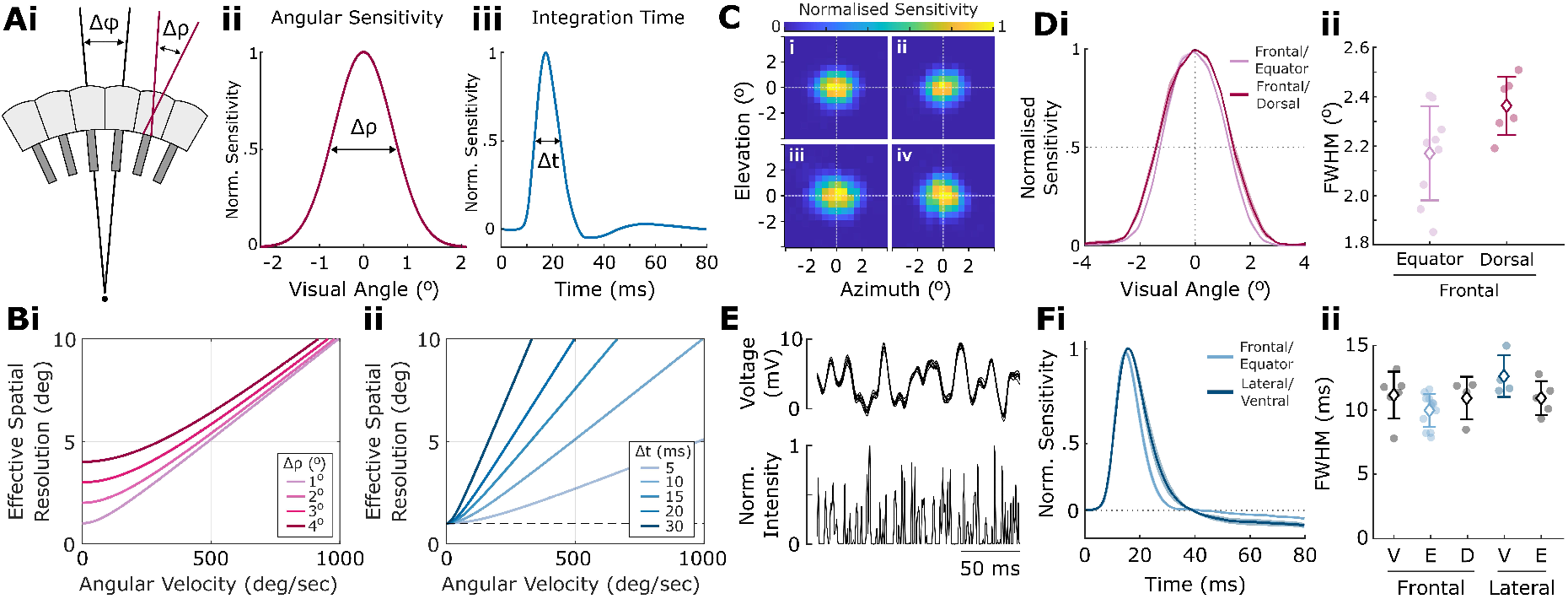
Photoreceptor Spatial and Temporal Resolution in Monarch Butterflies. **(A)** (i) Two anatomical features determine the maximum spatial visual acuity of the insect compound eye: the inter-ommatidial angle between adjacent facets, Δφ, which specifies the Nyquist sampling frequency of the eye lattice by 1/2Δφ; and the angular sensitivity function of the rhabdom, characterised by the full width at the sensitivity half-maximum, Δρ (ii). Δρ specifies the spatial cut-off frequency of each individual rhabdom as 1/Δρ. (iii) Impulse response of a light-adapted butterfly photoreceptor, adapted from Rutowski and Warrant 2002. The finite temporal resolution of phototransduction confers a delay followed by voltage response characterised by the full width at the sensitivity half-maximum, Δt. **(B)** Effective visual acuity, calculated using Snyder’s approximation ^7^. (i) Effective visual acuity as a function of sensor rotational velocity shown for rhabdom angular sensitivity angles Δρ ranging from 1°-4°. (ii) Effective visual acuity as a function of sensor rotational velocity shown for photoreceptor integration times Δt ranging from 5 to 30 ms. **(C)** Photoreceptor receptive fields in the (i-ii) frontal and (iii-iv) dorsal visual field. n = 4 cells from 1 animal. **(D)** (i) Angular acceptance sensitivity profiles of photoreceptors sampling the frontal (light pink) and dorsal (dark red) visual field. Frontal, n = 9 cells 2 animals; Dorsal, n = 6 cells, 2 animals; mean ± SEM. (ii) Full width at half-maximum (FWHM) of the cells in (i), corresponding to Δρ. **(E)** Photoreceptor temporal impulse responses were estimated by reverse correlation of photoreceptor voltage (top trace) with clipped white noise intensity fluctuations (bottom trace). Example shows an overlay of photoreceptor responses to multiple stimuli repetitions. **(F)** (i) Normalised temporal impulse responses of photoreceptors sampling the frontal equatorial (light blue; n = 12 cells, 4 animals) and ventrolateral (dark blue; n = 4 cells, 1 animal) visual field. Mean ± SEM. (ii) Full width at half-maximum (FWHM) of photoreceptor temporal impulse responses (corresponding to integration time, Δt) at separate locations across the visual field. V: ventral; E: equatorial; D: dorsal.

Here, Δρ is the quiescent photoreceptor angular acceptance width, and Δ*t* is the photoreceptor integration time. As shown in Figure 3B, *Δρ_eff_* increases with angular velocity, approaching a linear relationship with slope *Δt* at higher angular velocities.

To calculate *Δρ_eff_* following motion blur, we first measured photoreceptor Δρ and Δt using intracellular electrophysiology from quiescent light-adapted butterflies (Figure 3C-D). To estimate Δρ, we measured photoreceptor receptive fields from different locations across the eye (Figure 3C), defining Δρ as the full width at half-maximum (FWHM) of the receptive-field^37^. Photoreceptor Δρ varied across the eye as described previously^37^, with Δρ = 2.17 ± 0.19° in the frontal visual field (mean ± SD, n = 9 cells, 2 animals), broadening to 2.36 ± 0.12° in the dorsal visual field (n = 6 cells, 2 animals; Figure 3D). To estimate Δt, we calculated the photoreceptor temporal impulse response by reverse-correlating membrane voltage with clipped white-noise intensity fluctuations (Figure 3E). Photoreceptor Δt also varied across different eye regions (Figure 3F). Δt was narrowest in the frontal-equatorial visual field (9.97 ± 1.28 ms, mean ± SD, n = 12 cells, 4 animals), and largest in the lateral-ventral visual field (12.62 ± 1.62 ms, n = 4 cells, 1 animal).

Based on these electrophysiological measurements of Δρ and Δt, we next estimated the effective angular acceptance of photoreceptors during take-off flight and assessed the extent to which gaze stabilisation limits motion-induced degradation of spatial acuity (Figure 4). We first used the reconstructed 6-DOF head kinematics to calculate the optic-flow field across the visual sphere arising from the rotational and translational components of self-motion, and repeated the calculation using thorax kinematics to provide an unstabilised reference. At each position in visual space, we calculated the magnitude of the optic-flow vector tangent to the visual sphere (Figure 4A), corresponding to the absolute angular speed of retinal image motion, *ω*. Taking the median across time produced the characteristic angular speed seen at each location in the visual field (Figure 4Aii). Combining these angular speed distributions with photoreceptor Δρ and Δt using Equation (1) produces the effective photoreceptor acceptance angle, *Δρ_eff_*.

**Figure 4:**
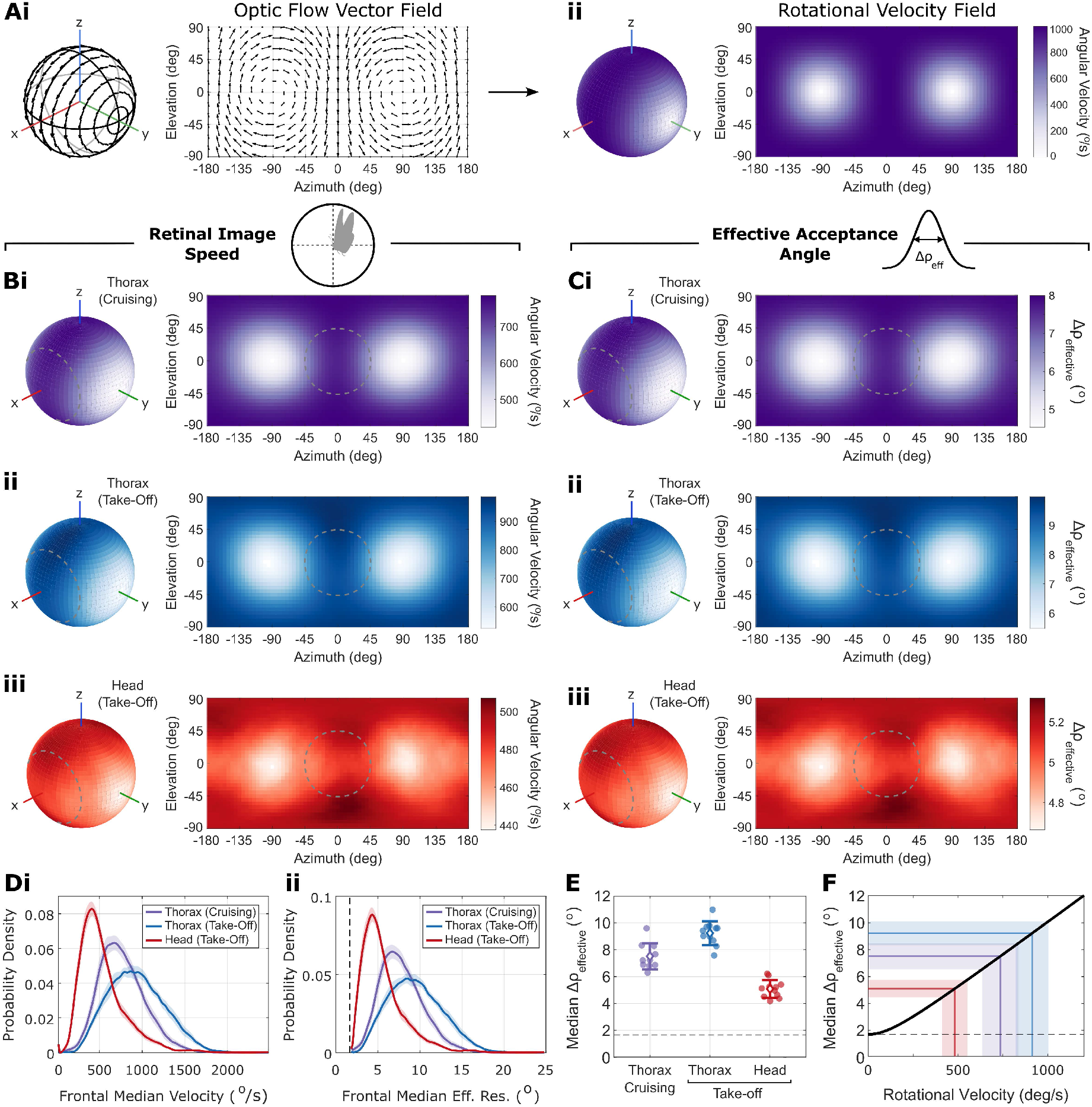
Motion Blur and the Effective Visual Acuity of Monarch Butterflies in Flight. **(A)** (i) Example optic flow snapshot for a pure pitch rotation. Vectors are tangent to the animal’s visual sphere and represent the direction and angular speed of local retinal image motion resulting from self-motion. Left, flow field in spherical coordinates relative to the animal’s frame of reference, where x = frontal, y = left, z = dorsal. Right, the same flow field as a cylindrical projection, where (0,0) corresponds to the animal’s frontal axis. For pure pitch rotation, angular image speed is minimal at the rotation axis passing through (−90,0) and (90,0). (ii) The magnitude of the optic-flow vector tangent to the visual sphere defines the angular speed of retinal image motion, ω. The angular distance travelled across the retina during photoreceptor integration is given by ωΔt. **(B)** Spatial distributions of retinal image speed calculated from 6-DOF kinematics for (i) thorax motion during cruising flight, (ii) thorax motion during take-off, and (iii) head motion during take-off. Thorax-derived optic flow provides an unstabilised reference for comparison with the head during take-off. Optic flow was calculated at each time-point from the rotational and translational components of self-motion, assuming a uniform spherical visual environment. At each azimuth and elevation, the magnitude of the optic-flow vector tangent to the visual sphere was calculated, and the median taken over time for each spatial location. Maps show the mean of these temporal medians across trajectories at each location in the visual field (cruising: n = 50 trajectories from 10 animals; take-off: n = 43 trajectories from 11 animals). Dashed grey circle shows the frontal visual area used for quantification in (D-F). **(C)** Spatial distributions of effective photoreceptor angular acceptance width, *Δρ_eff_*, calculated by combining the retinal image speed distributions in (B) with Δρ and Δt measured from intracellular electrophysiology according to Equation (1). Mean frontal visual-field values of Δρ = 2.17° and Δt = 9.97 ms were used. Dashed grey circle indicates the frontal visual-field region used for quantification in (D-F). **(D)** (i) Distribution of rotational image speeds within the frontal visual-field region indicated by the dashed grey circle in (B–C), calculated from thorax motion during cruising flight (purple), thorax motion during take-off (blue), and head motion during take-off (red). (ii) Corresponding distributions of effective photoreceptor acceptance angle, *Δρ_eff_*, within the same frontal visual region. **(E)** Median effective photoreceptor acceptance angle, *Δρ_eff_*, for the distributions shown in (Dii). Points show median values per animal; diamonds and error bars show mean ± SD across animals. **(F)** Relationship between retinal image speed and effective photoreceptor acceptance angle, *Δρ_eff_*, calculated from Equation (1) using the mean frontal visual-field values Δρ = 2.17° and Δt = 9.97 ms measured using intracellular electrophysiology (solid black line). Dashed horizontal line shows the quiescent acceptance angle, Δρ = 2.17°. Colored lines and shaded regions show the mean ± SD across animals of median *Δρ_eff_* values for thorax motion during cruising flight (purple), thorax motion during take-off (blue), and head motion during take-off (red).

Angular speed distributions for the thorax (Figure 4Bi-ii) reflected the prominent pitching oscillations, with minimal angular speeds occurring at the lateral poles of the pitching (y) axis, and average speeds in the frontal visual field of 737.3 ± 94.6 °·s⁻¹ during cruising flight, and 915.4 ± 105.3 °·s⁻¹ during take-off. Similarly, the average angular speed distributions for the head revealed a pitching component (Figure 4Biii), but this was spread more broadly across the visual field, with a slower average speed of 482.3 ± 70.0 °·s⁻¹ in the frontal visual field. These angular speed distributions corresponded to an average acceptance width of 5.28 ± 0.64° for the head in the frontal visual field. This compares to average acceptance widths of 7.67 ± 0.91° and 9.38 ± 1.02° for the unstabilised thorax in cruising and take-off flight, respectively. Gaze stabilisation therefore reduces effective acceptance width by 31.2% relative to the thorax during cruising flight and by 43.7% during take-off flight (Figure 4E), but nonetheless remained 2.4-fold broader than the anatomical Δρ (Figure 4F).

Interestingly, most of the retinal image motion experienced in flight arose from rotational rather than translational optic flow. Since translational optic flow depends on viewing distance, we initially assumed a uniform spherical environment with all visual features located 1 m from the animal. Under this world model, median angular image speeds in the frontal visual field arising from rotational optic flow were 730.5 ± 94.5 °·s⁻¹ for thorax motion during cruising flight, 903.7 ± 101.3 °·s⁻¹ for thorax motion during take-off, and 474.7 ± 68.6 °·s⁻¹ for head motion during take-off (mean ± SD). By comparison, the corresponding angular image speeds arising from translational optic flow were substantially lower at 52.9 ± 14.7 °·s⁻¹, 33.9 ± 6.1 °·s⁻¹, and 34.3 ± 5.9 °·s⁻¹, respectively. Under a non-uniform world model with a ground distance of 0.5 m, these translational components increased only slightly, to 55.1 ± 15.2 °·s⁻¹, 35.2 ± 6.2 °·s⁻¹, and 36.3 ± 6.0 °·s⁻¹, respectively.

## Discussion

Here we report estimates of spatial visual acuity in freely flying Monarch butterflies that account for the effects of motion blur^7,9^. We show that compensatory head movements substantially attenuate the large thoracic oscillations generated during flight, but do not fully stabilise the retinal image. Thoracic motion was dominated by wingbeat-coupled pitching oscillations (Figure 1), which were strongly attenuated at the head (Figure 2). Despite this compensation, substantial retinal image motion remained, broadening the effective photoreceptor acceptance angle (Δρ) from an anatomical value of 2.17° (Figure 3) to 5.28° during take-off (Figure 4). Whilst this represents a substantial improvement in image quality compared with the predicted effective acceptance angle of 9.38° for an unstabilised head, retinal image motion nevertheless degraded spatial resolution considerably relative to the anatomical limit.

Importantly, this broadening of effective photoreceptor acceptance angle does not represent a hard limit for object detection. High-contrast objects can remain detectable even when subtending well below the photoreceptor acceptance angle, provided they generate sufficient photoreceptor signal against the background^38,39^. Indeed, freely flying *Papilio xuthus* are able to detect colored disks subtending angles as small as approximately half their photoreceptor acceptance width^40^. However, this ability to detect isolated objects differs from the capacity to resolve spatial structure. The photoreceptor acceptance angle, Δρ, determines the degree of spatial low-pass filtering^7^, with a characteristic cut-off frequency of *v_co_* = 1/*Δρ*. Broadening of the photoreceptor acceptance function therefore progressively attenuates high spatial frequencies, degrading the spatial information available for distinguishing closely spaced objects and resolving fine visual patterns.

The spatial information transmitted by individual photoreceptors is in turn sampled by the ommatidial lattice to form an image, with the highest resolvable spatial frequency determined by the interommatidial angle, Δφ. The Nyquist sampling frequency, *v_s_*, of the compound eye lattice is approximately *v_s_* = 1/(2*Δφ*) along ommatidia rows, and *v_s_* = 1/(√3*Δφ*) along the diagonal^7^. For the frontal eye, our measured quiescent Δρ = 2.17° corresponds to a photoreceptor cut-off frequency of 0.46 cycles·deg^-1^. Assuming an interommatidial angle of Δφ = 1.5° (within the range of previous anatomical measurements^41^), gives a row-wise lattice Nyquist frequency of ∼0.33 cycles·deg^-1^. Stationary photoreceptors therefore transmit spatial frequencies substantially higher than those that can be sampled by the ommatidial lattice.

Spatial filtering and sampling are matched when *v_co_* ≈ *v_s_*, corresponding to Δρ/Δφ ratios of 1.7–2. Previous anatomical measurements in a variety of insect species report Δρ/Δφ ratios in the range 1-1.6, corresponding to an under-sampled regime in which individual photoreceptors transmit more spatial information than can be sampled by the ommatidial lattice^8,37^. This excess photoreceptor bandwidth has been proposed to provide a margin for motion blur during flight, where broadening of Δρ by retinal image motion progressively attenuates high spatial frequencies, shifting the effective Δρ/Δφ ratio towards the matched-sampling condition^8^. Our assumed frontal Δφ of 1.5° gives Δρ/Δφ = 1.45 for the quiescent Monarch eye, within the typical under-sampling range of insect eyes.

Our measurements directly quantify this effect during free flight and show that retinal image motion during take-off is sufficient to carry the eye beyond the matched condition. Motion blur broadened the effective acceptance angle to 5.28°, increasing Δρ_eff_/Δφ to 3.52 and reducing the effective photoreceptor cut-off frequency from approximately 0.461 (quiescent) to 0.189 cycles·deg^-1^. Without compensatory head movements, the predicted Δρ_eff_ of 9.38° would correspond to a Δρ_eff_/Δφ of 6.25 and a cut-off frequency of only 0.107 cycles·deg^-1^. Gaze stabilisation therefore increases the effective spatial cut-off by approximately 78% relative to the unstabilised condition, substantially increasing the spatial bandwidth available during take-off. Nevertheless, the gaze-stabilised cut-off remains below the row-wise lattice Nyquist frequency of approximately 0.33 cycles·deg^-1^, indicating that the eye remains motion-limited during take-off.

This lattice over-sampling may reflect the unusually large thoracic rotational velocities experienced during take-off. Take-off requires rapid generation of sufficient aerodynamic force to produce lift, and thoracic rotations are thought to help redirect the aerodynamic force vector downwards^34^. During cruising flight, we found that thoracic oscillations are slower and of smaller amplitude than during take-off (Figure 1M-O). Assuming the same proportional attenuation of thorax oscillation during cruising flight as measured during take-off (head/thorax angular-velocity ratio = 0.53), the lower thoracic rotational velocities predict a gaze-stabilised effective acceptance angle of approximately 4.46°, corresponding to a Δρ_eff_ /Δφ ratio of 2.97. Thus, under less demanding flight conditions, gaze stabilisation may bring spatial sampling closer to, although still beyond, the matched regime.

Interestingly, butterflies make extensive use of gliding during flight, with many species alternating bouts of active flapping with periods of gliding^27,42–44^. Thoracic oscillations are likely to be substantially reduced during gliding, thereby further reducing rotational retinal image motion following gaze stabilisation. The resulting reduction in motion blur would shift Δρ_eff_/Δφ further towards the matched-sampling condition. It will therefore be interesting to determine to what extent gliding provides periods of enhanced spatial visual performance, and whether butterflies actively exploit these periods during behaviours that demand heightened visual acuity, such as detecting conspecifics^42,43^ or food sources.

Furthermore, whilst our analyses here primarily characterise the average optic flow experienced during free flight, the rotational velocities underlying this optic flow vary continuously across the wingbeat cycle. Wingbeat-coupled thoracic oscillations contain phases of high rotational velocity interspersed with phases in which rotational velocity approaches zero, with corresponding phase-dependent changes in head velocity and retinal image motion. These brief periods of reduced retinal image velocity may provide opportunities to extract fine spatial information that would otherwise be attenuated by motion blur. Such temporal variation in spatial acuity could contribute to the ability of *Papilio xuthus* to detect small chromatic targets^40^. This raises the possibility that butterflies preferentially extract fine spatial information during periods of reduced retinal image motion, with visual discrimination performance varying systematically across the wingbeat cycle.

Overall, our results show the significant demands placed on the neck motor system of flying butterflies. The large thoracic oscillations inherent to lepidopteran flight must be compensated by movements of the head, otherwise motion blur would substantially degrade visual acuity. Whilst gaze stabilisation reduces the loss of spatial information in flight, the effective photoreceptor acceptance angle remains more than twice its quiescent width and exceeds the matched-sampling regime of the ommatidial lattice. Gaze stabilisation therefore does not fully eliminate motion blur, highlighting how locomotor dynamics shape the visual performance of an animal, and, ultimately, its sensory ecology.

## Methods

### Animals

Monarch butterflies (*Danaus plexippus*, RRID: NCBITaxon_13037) were purchased as pupae from Stratford Butterfly Farm (UK) and The Entomologist Ltd. (UK). Pupae were stored at 23°C and 60% humidity on a 12-hr dark/light cycle. Imagoes were left to harden for 24-hrs post-emergence, then stored at 15°C. Butterflies were fed dilute honey every other day, and all experiments were performed within two weeks post-emergence.

### Motion-Capture Reconstruction of Thorax Kinematics

Free-flight motion capture trajectories were acquired in a cylindrical indoor flight enclosure measuring 2.0 m diameter and 2.4 m high, made from a matte diffusing white fabric (White Joelastic fabric, J & C Joel, UK). A panoramic naturalistic scene of trees and foliage covered the lower 1m of the arena to provide visual contrast and horizon. The enclosure was illuminated by four CDM-TMW Elite 315 W broad-spectrum HID lamps (Philips, Netherlands), diffused through the enclosure ceiling. Butterflies were gently handled by the wings and dropped by hand from approximately 1m from the ground.

Thorax kinematics were measured by tracking three points on a small T-shaped carbon-fibre frame affixed to the meso-metathoracic notum such that the wings did not clash with the marker (Figure S1). Cruising-flight kinematics were measured using eight infrared (850 nm) motion-capture cameras at 180 fps (Miqus M5, Qualisys AB, Sweden). Three 1 mm diameter resin-printed spheres were covered in retro-reflective tape (product no. 160054, Qualisys AB, Sweden) and affixed to the carbon-fibre T-frame with UV glue (Figure S1). Individual marker trajectories were automatically tracked and manually labelled using Qualisys Track Manager software (Qualisys AB, Sweden). Reconstructed inter-marker distances had a mean within-recording standard deviation of 0.16 mm (N=29 recordings; Figure S1F).

Thorax orientation and centre-of-mass were estimated from reconstructed motion-capture marker positions using a marker-to-thorax transformation calculated prior to each recording. Stereo photographs were used to 3D reconstruct the three marker centres and six naturally occurring white spots in the monarch’s thoracic patterning (Figure S1C). Three spots on each side of the thorax were used to define the thorax coordinate system, from which the rigid rotation and translation relating the marker frame to the thorax frame were calculated (Figure S1DE). This stereo photograph method aligned closely with measurements taken from a high-resolution photogrammetric scan (Figure S1E).

Motion-capture T-frames weighed 10.0 ± 0.9 mg. To assess the potential impact of marker frame placement on flight kinematics, we tested whether marker orientation and position predicted thorax roll, pitch, or yaw rotational velocities (Figure S2). Variation in marker placement was not associated with bias in rotational velocity within animals or across pooled recordings (linear mixed-effects models accounting for individual differences; all *p* > 0.094 before adjusting for multiple comparisons; 29 marker placements across 10 animals).

### Multi-Camera Videographic Reconstruction of Head and Thorax Kinematics

Take-off kinematics were measured in the same flight enclosure using multi-camera high-speed videography at 1000 fps (two FASTCAM SA3, Photron Ltd., Japan; two Chronos 1.4, Kron Technologies, Canada; four AF Nikkor 50 mm f/1.8D lenses, Nikon, Japan; Figure S1G) and digitised at 500 fps. Butterflies were placed on a small branch approximately 50 cm from the ground and left to take off spontaneously. Camera acquisition was end-triggered simultaneously from a manual button-press, and frame-synchronised post-hoc from a small red LED flashing within the field of view of all cameras.

Thorax kinematics were measured from a carbon fibre T-frame affixed to the meso-metathoracic notum as for motion-capture, but the three tracking points were instead painted with white nail varnish (Figure S1H). These T-frames weighed 2.1 ± 0.2 mg. T-frame tracking points were digitised using DeepLabCut^45^ and reconstructed using Anipose^46^. Across all recordings, reconstructed T-frame lengths had a mean error of 0.053 ± 0.062 mm (mean ± SD; N = 43 recordings; Figure S1I). Thorax orientation and centre-of-mass were estimated using a marker-to-thorax transformation for each unique marker placement. The thorax coordinate frame was defined from a triangle formed between two white dots on the tegulae (i.e. a mesothoracic sclerite covering the dorsal forewing hinge) and the dorsal tip of the mesoscutellum at maximum downstroke (Figure 1A).

Head kinematics were reconstructed from each frame using at least three out of five naturally occurring white spots on the vertex of the head cuticle (Figure 1A, S1J). Head points were digitised manually and reconstructed using Anipose^46^. The head coordinate system for each video frame was estimated using a white-spot-to-head transformation defined from a high-resolution photogrammetric scan (Figure S1J). The lateral head (y) axis was defined from the normalised mean vector between left and right white head spots. A provisional dorsal (z) axis was defined following Rutowski and Warrant, 2002, from the dorsoventral orientation of the slightly flattened posterior aspect of each eye^37^ (Figure S1Jii). The frontal (x) axis was defined as the vector normal to the y and provisional z axes. The final z axis was then orthogonalized relative to the x-y plane. The head centre was defined as the intersection of the line connecting the centres of the left and right eyes and the x-z plane. The accuracy and precision of head tracking were evaluated relative to a ground-truth reference (Figure S1K-O). Pitch orientation estimates derived from the head spots had a mean error of −0.026 ± 1.09° (mean ± SD) for the raw data, whereas smoothing the xyz coordinates reduced this to −0.020 ± 0.85° (Figure S1N).

### Photoreceptor Intracellular Recordings

The recordings were performed as described previously^47^. Briefly, the butterflies were immobilized with bees wax and resin (3:1) with the eye positioned in the rotation centre of a dual goniometric and x-y-z stage, carrying the micromanipulator (Sensapex, Finland) and the preamplifier (NPI, Germany). A microelectrode (100-150 MΩ resistance), produced from 1/0.5 mm outer/inner diameter borosilicate capillaries with a laser puller (P-2000, Sutter, USA), back-filled with 3M KCl, was inserted into the compound eye through a small hole in the cornea, cut with a razorblade chip. Impaled photoreceptors were quickly analysed with an array of monochromatic LEDs^48^; cells, peaking around 540-580 nm and without any color-opponent negative responses^49^ were considered as the photoreceptors R3-8 and analysed further.

The photoreceptor receptive field was mapped by measuring the voltage response to a 0.5×0.5° white square on a 10×10° virtual grid, centered on the photoreceptor’s optical axis, back-projected on a glass diffuser from an RGB DLP projector (LightCrafter 4500, Texas Instruments, USA) with a high frame rate (220 Hz), using PsychoPy software package^50^. The photoreceptor impulse responses were measured by stimulating the impaled photoreceptors with repeated pseudorandom stimuli, i.e. modified Gaussian white-noise stimulation, delivered with a linearly driven white LED (OptoLED, Cairn, UK). The LED intensity time series, whitened between 1 and 500 Hz, was generated in Matlab; the negative values were clipped to 0 for increased contrast^51^.

## Data Analysis

### Flight trajectory analysis

Coordinate systems for the head and thorax defined x-, y-, and z-axes as frontal, lateral, and dorsal, respectively. Roll, pitch, and yaw rotations were defined about these x-, y-, and z-axes. Positive roll corresponded to elevation of the left side of the thorax, positive pitch to nose-up rotation, and positive yaw to rightward rotation. Relative orientations were obtained from rotation matrices using a Z–Y–X Euler-angle decomposition, corresponding to sequential yaw, pitch, and roll components. Angular velocities were calculated from changes in the body-axis vectors using second-order central differences after cubic-spline interpolation to 1-ms intervals, with incremental rotations treated under a small-angle approximation. Angular accelerations were calculated as the temporal derivative of the corresponding angular velocity traces.

To identify the dominant axis of rotational motion, principal component analysis (PCA) was performed separately for each flight trajectory. The first principal component (PC1) described the axis in roll–pitch–yaw angular-velocity space accounting for the greatest variance in rotational motion. Because the sign of principal components is arbitrary, PC1 orientation was standardised by defining the positive direction as that with a positive pitch component.

Kinematic distributions are presented as probability densities. For each animal, all trajectory time points were pooled to generate an animal-level distribution, and the mean ± SEM across animal-level distributions was plotted. These mean distributions were summarised using the median and interquartile range (IQR), as indicated in the main text.

### Kinematic data smoothing and derivative estimates

Kinematic data were filtered with a forward-reverse (zero-phase) 3^rd^ order low-pass Butterworth filter. Estimating velocity and acceleration from kinematic data is inherently sensitive to noise, as differentiation amplifies high-frequency fluctuations. Furthermore, excessive smoothing can conversely underestimate the true derivative. To balance noise reduction against signal preservation, an optimal low-pass filter cut-off frequency was chosen to smooth the data, following the method described in^52^ Bomphrey et al., 2009 (Figure S3). This method assumes that the underlying kinematic signal is temporally correlated, whereas measurement noise is temporally uncorrelated. Under this assumption, the auto-correlogram (ACG) of residuals obtained by subtracting the filtered signal from the raw data will differ depending on whether those residuals contain primarily noise or retained signal (Figure S3A). If the residuals consist mainly of noise, the ACG is expected to be largely flat, with only a sharp peak at zero lag, and therefore to have low variance. In contrast, if residuals still contain true signal, the ACG will show a broader peak that decays towards zero with increasing lag, resulting in higher variance. The optimal low-pass cut-off frequency was therefore defined as the lowest cut-off frequency at which residual ACG variance was minimised (Figure S3Aiii), indicating that the residuals consisted predominantly of uncorrelated noise and contained minimal true signal.

Estimated optimal low-pass cut-off frequencies for raw xyz data for were 48 Hz for motion capture T-marker, 65 Hz for take-off T-marker, and 60 Hz for take-off head points (Figure S3D-F). Optimal low-pass cut-off frequencies were also calculated for rotational velocities (60/100/100 Hz), and rotational accelerations (68/130/140 Hz) for motion capture T-marker, take-off T-marker, and take-off head points, respectively (Figure S3D-F). Optimal filters for the head reconstruction were first evaluated relative to a ground truth reference triangle (Figure S1K-O). Raw head angular velocities were substantially higher than those of the reference triangle (median: 220.1 vs. 61.3 deg/s, interquartile range, IQR: 108.3-373.5 deg/s vs 30.7-105.5 deg/s; raw head and reference triangle, respectively; Figure S1O). Smoothing reduced the median head angular velocity to 69.3 deg/s, with an interquartile range of 31.0-113.7 deg/s, closely matching the reference triangle distribution (Figure S1O). The effect of low-pass filtering on derivative estimates was further evaluated by comparing derivative histograms for a range of low-pass cut-off frequencies (Figure S4).

### Photoreceptor electrophysiology analysis

The voltage responses to the pattern stimulus (flashing square on a virtual grid) were transformed into sensitivities according to the stimulus-response curve, fitted with a Hill sigmoid function^48^. A two-dimensional Gaussian was fitted to each receptive field. Angular full widths at half maximum (FWHM) were calculated along the two principal axes of the fitted Gaussian. Photoreceptor acceptance angle, Δρ, was defined as the geometric mean of the major-and minor-axis FWHMs.

The photoreceptor impulse response was determined from the responses to repeated pseudorandom Gaussian white noise stimuli. For each recording, the frequency-dependent transfer function between the stimulus frequency spectrum and the response frequency spectrum was calculated in MATLAB and averaged across repetitions. The time-domain impulse response was then obtained by inverse Fourier transformation of the transfer function. Integration time, Δt, was defined as the full width at half maximum (FWHM) of the temporal impulse response, measured relative to the response baseline.

## Acknowledgements

Thanks to Oscar Healy for assistance with acquiring high-resolution photogrammetric scans. This work was supported by Air Force Office of Scientific Research (AFOSR) grant FA8655-23-1-7049 to GB and HGK, Biotechnology and Biological Sciences Research Council (BBSRC) grant BB/X002276/1 to HGK. Thanks to Ric Wehling and Nandini Iyer for providing financial support.

## Author Contributions

Conceptualization, JAS; Methodology, JAS/WL/MI/MK/HTL/GB; Software, JAS/MI/WL; Formal Analysis, JAS/WL; Investigation, JAS/WL; Resources, HTL/GB/HGK; Writing – Original Draft, JAS; Writing – Review & Editing, JAS/WL/MI/MK/HTL/GB/HGK; Visualization, JAS/WL; Supervision, HGK; Project Administration, JAS/HGK/GB; Funding Acquisition, JAS/HTL/GB/HGK.

---

**Figure S1:**
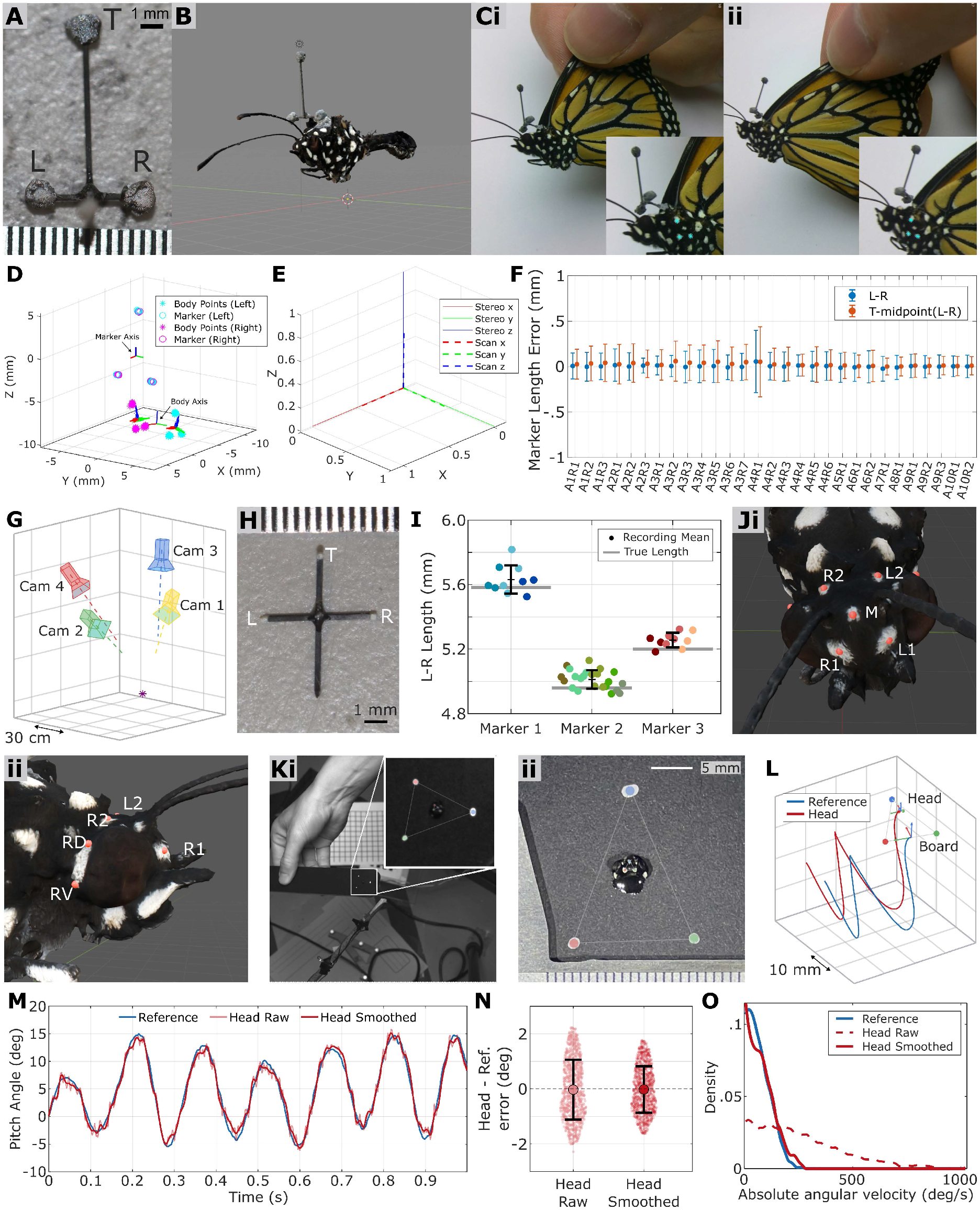
Motion-Capture and Videographic Kinematic Measurements. **(A)** Carbon fibre marker frame used for motion capture thorax kinematics. Ruler gradations are 0.5 mm. **(B)** High-resolution photogrammetric reconstruction of monarch butterfly body and marker for ground truth comparison. **(C)** Example (i) left and (ii) right stereo images used to estimate marker-to-thorax transformation matrices for each mounted marker prior to motion-capture recording. **(D)** 3D reconstructed T-marker points (circles) and body points (asterisks). Coordinate frame x/y/z (frontal/left/dorsal) shown in red/green/blue for the marker, body points on each side, and resultant body axis. **(E)** Comparison of motion-capture marker-to-thorax rotation matrices calculated using the stereo rig (dashed lines) compared to the high-resolution photogrammetric scan in B (solid lines). Note this compares the marker-to-thorax rotation matrices. x/y/z refer to frontal/left/dorsal. **(F)** Motion-capture marker length errors throughout each recording. Animals and recordings indicated in the form A#R#. **(G)** Example multi-camera arrangement for high-speed videography. **(H)** Example miniature carbon-fibre T-marker for multi-camera videographic thorax kinematics. **(I)** Ground-truth T-marker L-R lengths vs mean reconstructed lengths for multi-camera videography. Horizontal grey lines show ground-truth lengths for the three T-markers used for experiments. Circles show mean reconstructed lengths per recording, color-coded by animals. Black lines and error bars show mean ± SD across recordings for each T-marker. Across all recordings, reconstructed T-frame lengths had a mean error of 0.053 ± 0.062 mm (mean ± SD; n = 43 recordings). **(J)** Photogrammetric scan used to estimate head orientation from the five naturally occurring white spots that were tracked from videography. (i) Frontal view. Red dots show the white spots that were tracked, labelled Left (L), Right (R), and Medial (M). (ii) Side view. The posterior aspect of each eye is slightly flattened, which was used to define the dorsal axis of the eye, following Rutowski and Warrant 2002. Red dots show the two points (Right Dorsal, RD; Right Ventral, RV) used to define this axis. **(K)** (i) To validate the accuracy and precision of head coordinate frame reconstruction, we compared measured head orientation to a ground truth using a monarch butterfly head glued to a black piece of acrylic with three white points forming a reference triangle. (ii) The triangle formed by butterfly natural head spots L2-R2-M was compared with the reference triangle ground truth. **(L)** Example simultaneous tracking of the head and reference triangle shown in (K). **(M)** Normalised pitch orientation of the reference triangle (blue) vs head (red). Head orientation derived from raw reconstructed xyz points is shown (light red) along with that derived from smoothed head xyz points (dark red). The head was smoothened using the optimal filter described in Figure S4. **(N)** Orientation error between head and reference triangle. Raw head measurements had an error of −0.026 ± 1.09°, whilst smoothed head measurements had an error of −0.020 ± 0.85°. **(O)** Comparison of absolute angular velocity histograms for the reference triangle (blue line), raw head points (dashed red line), and smoothed head points (solid red line).

**Figure S2:**
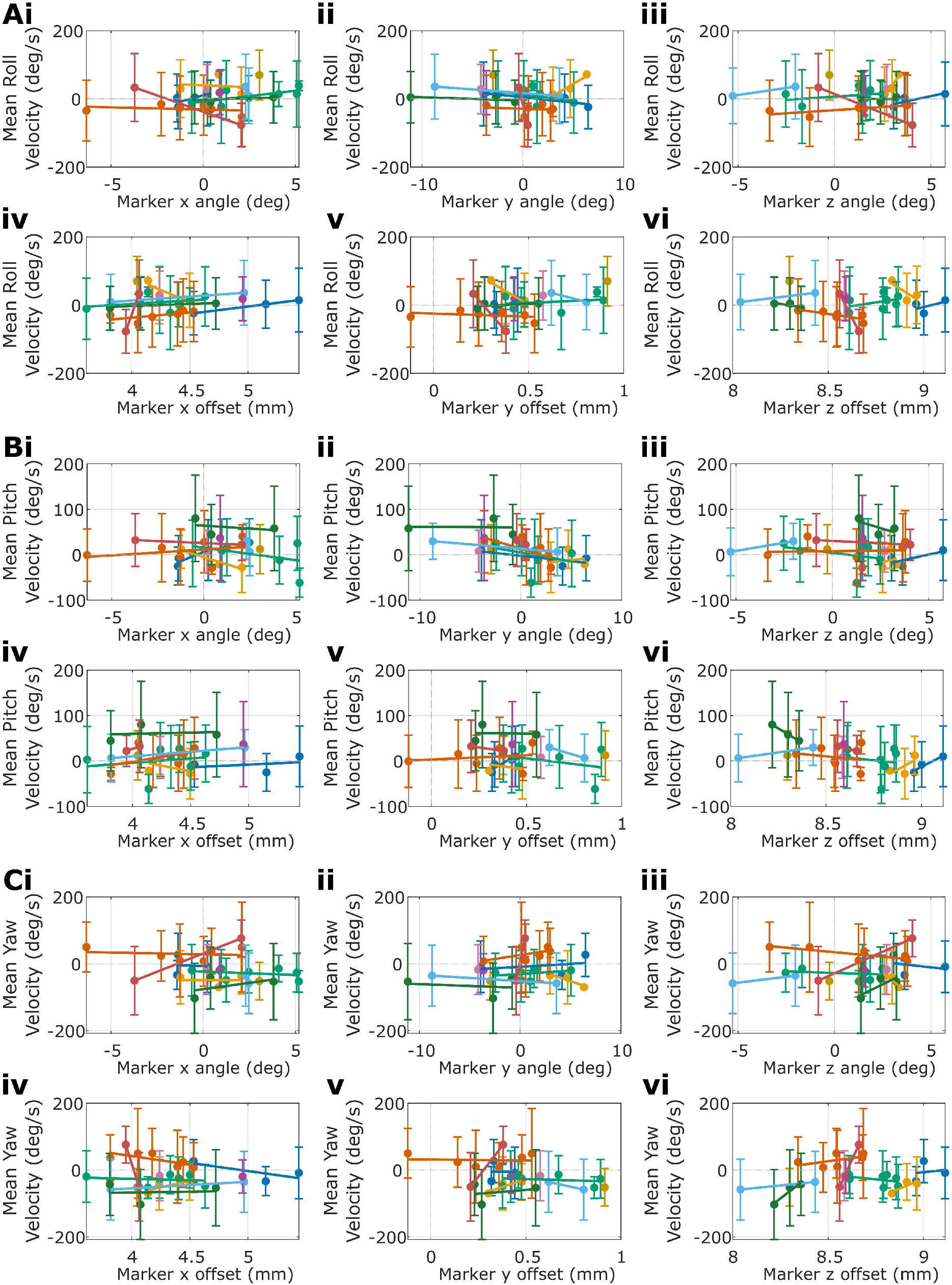
Relationship between marker placement and motion-capture free-flight measurements. **(A-C)** Motion-capture-derived thorax (A) roll, (B) pitch, and (C) yaw rotational velocities plotted against marker-frame (i-iii) orientation and (iv-vi) position relative to animal’s thorax coordinate frame. Each point shows the mean signed rotational velocity for one recording (marker position), with colours indicating individual animals (N=29 marker orientations across 10 animals). Error bars show the SD of per trajectory rotational velocity means within each recording. Coloured lines show within-animal linear regressions across recordings. (i) Marker x-axis rotational offset (approximately roll axis); (ii) Marker y-axis rotational offset (approximately pitch axis); (iii) Marker z-axis rotational offset (approximately yaw axis); (iv) Marker x-axis translation offset (approximately forward axis); (v) Marker y-axis translation offset (approximately left axis); (vi) Marker z-axis translation offset (approximately dorsal axis).

**Figure S3:**
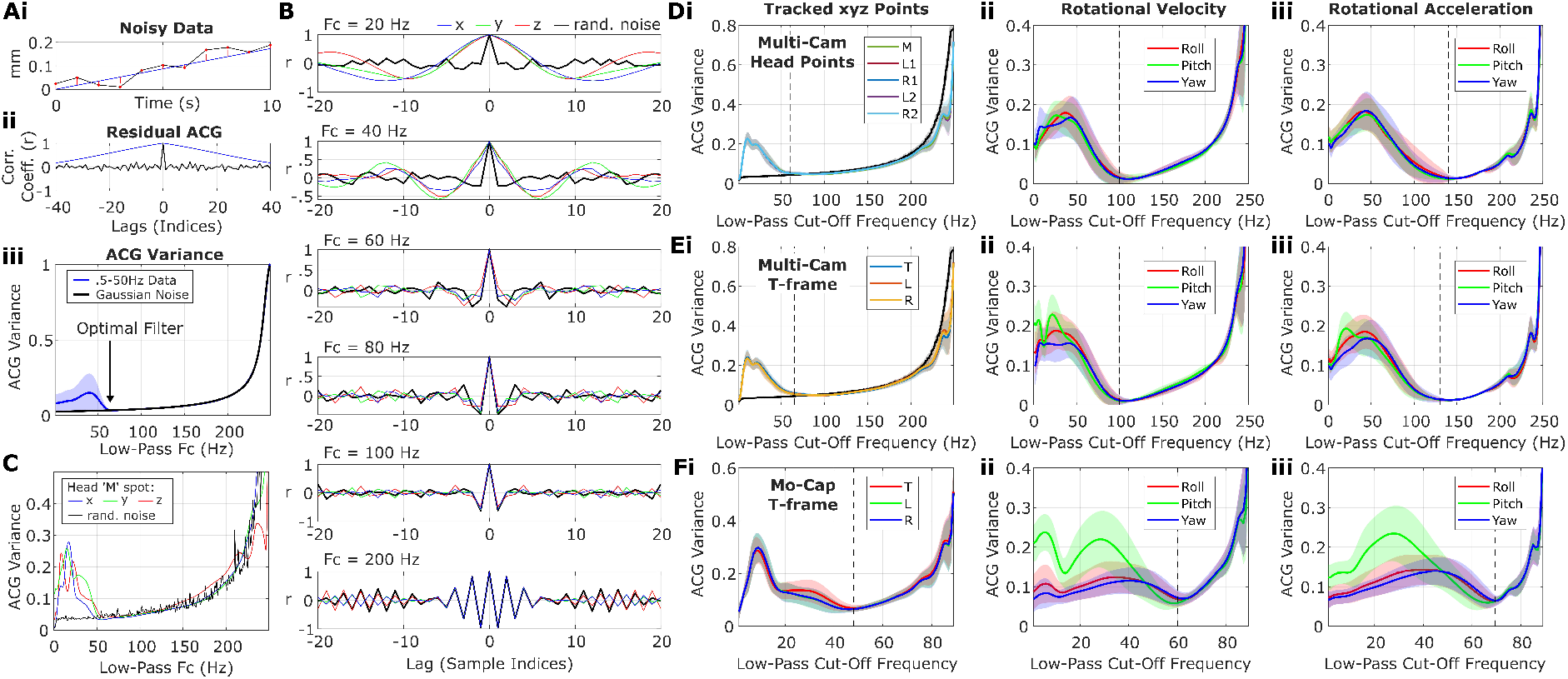
Optimal filter estimation for kinematic data. **(A)** Autocorrelation procedure to determine the optimal low-pass filter cut-off frequency for the data, based on ^52^. **(i)** Illustrative simulated data example (black line) of correlated signal (blue line) with additive noise (residuals in red). **(ii)** Auto-correlogram (ACG) of the additive noise residuals (black) and underlying signal (blue). Since the residuals are temporally uncorrelated, the residual ACG is mostly flat with a sharp peak at zero lag (i.e. overall low variance). In contrast, signal is typically correlated over time; as such, the signal ACG has a broad peak decaying to zero at large lags (i.e. overall high variance). **(iii)** Example ACG variance vs low-pass filter cut-off frequency for simulated 0.5-50 Hz signal with additive gaussian noise (blue), as compared with pure gaussian noise (black). ACG variance for the simulated data increases for low-pass filter cut-off frequencies below ∼60 Hz, as residuals contain increasingly more correlated signal. We aim to choose a filter that removes as much noise as possible without removing the signal. The optimal filter is specified as the smallest low-pass filter cut-off frequency where the residuals between raw and filtered data resemble noise (i.e., have a low variance ACG). Note that data is simulated at 500 Hz and ACG variance increases when filter cut-off frequency approaches the Nyquist frequency because the residuals approach 0. **(B)** Representative normalised ACGs for 3D-reconstructed butterfly middle (‘M’) head spot data with increasing low-pass filter cut-off frequencies (Fc). Red, green, and blue lines show the ACGs of residuals between raw and filtered x, y, and z data, respectively. Black lines show ACGs of random noise in the same sampling base for comparison. At low Fc (top panels), the data is over-smoothed, and x/y/z residuals contain correlated signal, producing broad, high-variance ACGs. As Fc is increased, smoothed data residual ACGs increasingly resemble noise. When Fc approaches the Nyquist frequency (bottom panel), the residuals between raw and filtered data approach zero and produce high variance ACGs. **(C)** ACG variance vs low-pass filter cut-off frequency for the 3D-reconstructed butterfly middle (‘M’) head spot data in B. Red/green/blue lines show the ACGs of residuals between raw and filtered x/y/z data. Black lines show ACGs of random noise in the same sampling base for comparison. **(D)** Filtered residual ACG variance vs low-pass filter cut-off frequency (Fc) for the multi-camera butterfly head reconstruction. (i) head spot xyz coordinates (averaged xyz for each head spot), (ii) rotational velocity, and (iii) rotational acceleration. Optimal cut-off frequencies in dashed black. **(E)** Filtered residual ACG variance vs low-pass filter cut-off frequency (Fc) for the multi-camera T-frame. (i) T-frame marker xyz coordinates (averaged xyz across each marker), (ii) rotational velocity, and (iii) rotational acceleration. Optimal cut-off frequencies in dashed black. **(F)** Filtered residual ACG variance vs low-pass filter cut-off frequency (Fc) for the motion-capture T-frame. (i) T-frame marker xyz coordinates (averaged xyz across each marker), (ii) rotational velocity, and (iii) rotational acceleration. Note, a sampling base of 180Hz was used for motion-capture. Optimal cut-off frequencies in dashed black.

**Figure S4:**
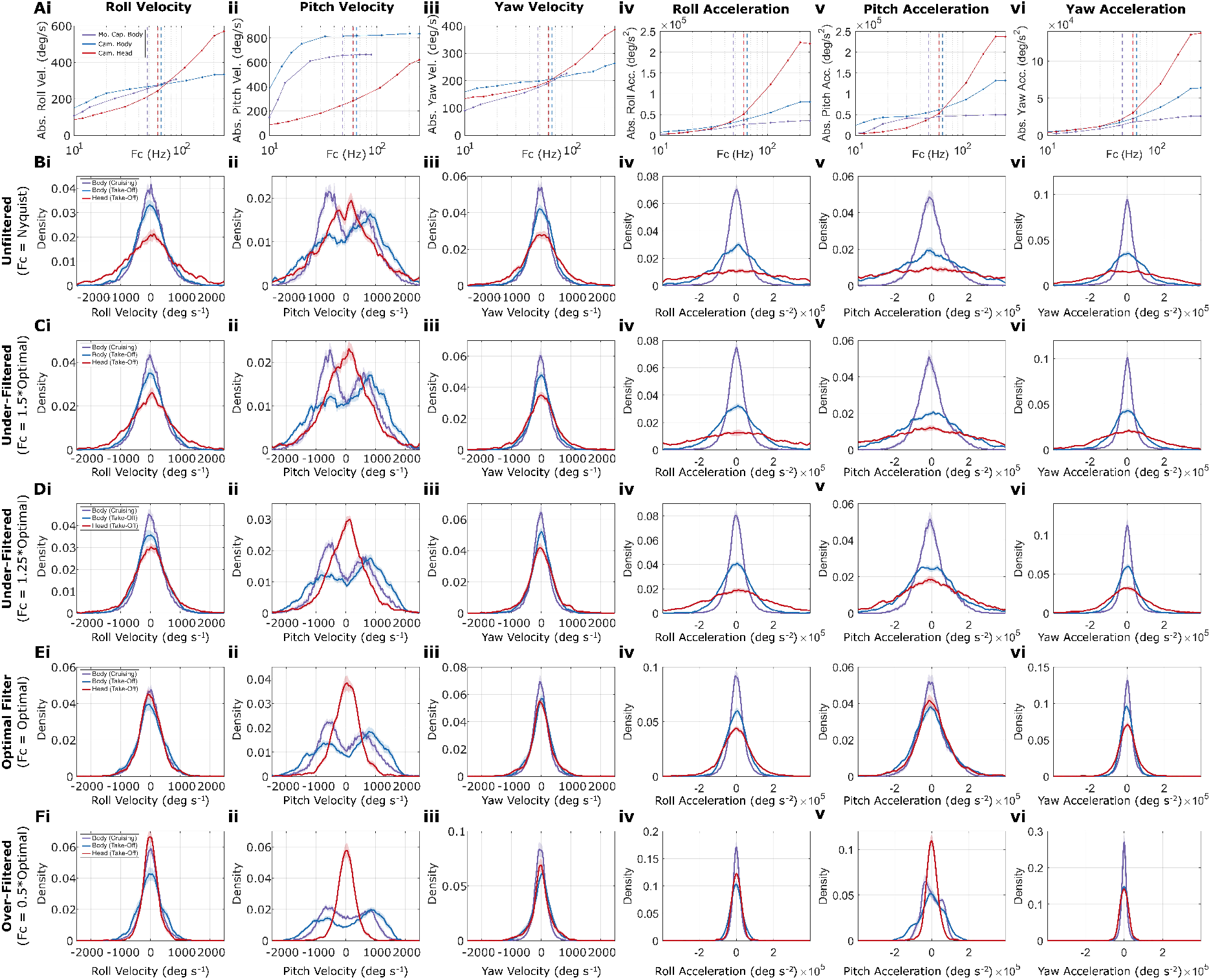
Effect of smoothing on kinematic derivative estimates. **(A)** Comparison of median absolute (i-iii) velocities and (iv-vi) accelerations across low-pass filter cut-off frequencies *F_c_*. Motion-capture (mo. cap.) thorax measurements are shown in purple, multi-camera (cam.) thorax measurements in blue, and multi-camera head-orientation measurements in red. Note that the Nyquist frequencies of the motion-capture and multi-camera data are 90 Hz and 250 Hz, respectively, which set the maximum possible *F_c_*. Dashed vertical lines show the optimal *F_c_* used for analysis. (i) roll velocity, (ii) pitch velocity, (iii) yaw velocity, (iv) roll acceleration, (v) pitch acceleration, and (vi) yaw acceleration. Legend in (i). **(B)** Unfiltered (i.e. raw data, or *F_c_* = Nyquist) velocity and acceleration histograms. Solid line and shaded area show mean ± SEM. Legend in (i). **(C)** Under-filtered velocity and acceleration histograms. *F_c_* = 1.5 · *Optimal F_c_*. **(D)** Under-filtered velocity and acceleration histograms. *F_c_* = 1.25 · *Optimal F_c_*. **(E)** Optimally filtered velocity and acceleration histograms. *F_c_* = *Optimal F_c_*. **(F)** Over-filtered velocity and acceleration histograms. *F_c_* = 0.5 · *Optimal F_c_*.

## References

1. Heras, F.J., and Laughlin, S.B. (2026). Investments in photoreceptors compete with investments in optics to determine eye design. eLife 13, RP96517. 10.7554/eLife.96517.

2. Laughlin, S.B., De Ruyter Van Steveninck, R.R., and Anderson, J.C. (1998). The metabolic cost of neural information. Nature Neuroscience 1, 36–41. 10.1038/236.

3. Okawa, H., Sampath, A.P., Laughlin, S.B., and Fain, G.L. (2008). ATP Consumption by Mammalian Rod Photoreceptors in Darkness and in Light. Current Biology 18, 1917–1921. 10.1016/j.cub.2008.10.029.

4. Pangršič, T., Stušek, P., Belušič, G., and Zupančič, G. (2005). Light dependence of oxygen consumption by blowfly eyes recorded with a magnetic diver balance. J Comp Physiol A 191, 75–84. 10.1007/s00359-004-0571-y.

5. Niven, J.E., Anderson, J.C., and Laughlin, S.B. (2007). Fly Photoreceptors Demonstrate Energy-Information Trade-Offs in Neural Coding. PLoS Biology 5, e116. 10.1371/journal.pbio.0050116.

6. Land, M.F., and Nilsson, D.-E. (2007). Animal Eyes (Oxford University Press).

7. Snyder, A.W. (1977). Acuity of compound eyes: Physical limitations and design. J. Comp. Physiol. 116, 161–182. 10.1007/BF00605401.

8. Land, M.F. (1997). Visual acuity in insects. Annual review of entomology. Vol. 42, 147–177. 10.1146/annurev.ento.42.1.147.

9. Srinivasan, M.V., and Bernard, G.D. (1975). The effect of motion on visual acuity of the compound eye: A theoretical analysis. Vision Research 15, 515–525. 10.1016/0042-6989(75)90029-2.

10. Snyder, A.W. (1979). Physics of Vision in Compound Eyes. In Comparative Physiology and Evolution of Vision in Invertebrates: A: Invertebrate Photoreceptors, H. Autrum, M. F. Bennett, B. Diehn, K. Hamdorf, M. Heisenberg, M. Järvilehto, P. Kunze, R. Menzel, W. H. Miller, A. W. Snyder, et al., eds. (Springer), pp. 225–313. 10.1007/978-3-642-66999-6_5.

11. Hardcastle, B.J., and Krapp, H.G. (2016). Evolution of Biological Image Stabilization. Current Biology 26, R1010–R1021. 10.1016/j.cub.2016.08.059.

12. Cellini, B., Salem, W., and Mongeau, J.-M. (2022). Complementary feedback control enables effective gaze stabilization in animals. Proceedings of the National Academy of Sciences 119, e2121660119. 10.1073/pnas.2121660119.

13. Hardcastle, B.J., Bierig, K., Heras, F.J., Longden, K.D., Schwyn, D.A., and Krapp, H.G. (2021). Passive inertial damping improves high-speed gaze stabilization in hoverflies. Preprint at bioRxiv, 10.1101/2021.12.11.472180.

14. Hengstenberg, R., Sandeman, D.C., and Hengstenberg, B. (1986). Compensatory head roll in the blowfly Calliphora during flight. Proc. R. Soc. B 227, 455–482. 10.1098/rspb.1986.0034.

15. Windsor, S.P., and Taylor, G.K. (2017). Head movements quadruple the range of speeds encoded by the insect motion vision system in hawkmoths. Proc. R. Soc. B 284, 20171622. 10.1098/rspb.2017.1622.

16. Cellini, B., and Mongeau, J.-M. (2022). Nested mechanosensory feedback actively damps visually guided head movements in Drosophila. eLife 11, e80880. 10.7554/eLife.80880.

17. Chatterjee, P., Prusty, A.D., Mohan, U., and Sane, S.P. (2022). Integration of visual and antennal mechanosensory feedback during head stabilization in hawkmoths. eLife 11, e78410. 10.7554/eLife.78410.

18. Mureli, S., Thanigaivelan, I., Schaffer, M.L., and Fox, J.L. (2017). Cross-modal influence of mechanosensory input on gaze responses to visual motion in Drosophila. J Exp Biol 220, 2218–2227. 10.1242/jeb.146282.

19. Boeddeker, N., Dittmar, L., Stürzl, W., and Egelhaaf, M. (2010). The fine structure of honeybee head and body yaw movements in a homing task. Proc. R. Soc. B 277, 1899–1906. 10.1098/rspb.2009.2326.

20. Boeddeker, N., and Hemmi, J.M. (2009). Visual gaze control during peering flight manoeuvres in honeybees. Proc. R. Soc. B 277, 1209–1217. 10.1098/rspb.2009.1928.

21. Doussot, C., Bertrand, O.J.N., and Egelhaaf, M. (2021). The Critical Role of Head Movements for Spatial Representation During Bumblebees Learning Flight. Front. Behav. Neurosci. 14. 10.3389/fnbeh.2020.606590.

22. Hateren, J.H.V., and Schilstra, C. (1999). Blowfly flight and optic flow: II. Head movements during flight. Journal of Experimental Biology 202, 1491–1500. 10.1242/jeb.202.11.1491.

23. Land, M.F. (1973). Head Movement of Flies during Visually Guided Flight. Nature 243, 299–300. 10.1038/243299a0.

24. Ravi, S., Siesenop, T., Bertrand, O.J., Li, L., Doussot, C., Fisher, A., Warren, W.H., and Egelhaaf, M. (2022). Bumblebees display characteristics of active vision during robust obstacle avoidance flight. Journal of Experimental Biology 225, jeb243021. 10.1242/jeb.243021.

25. Riabinina, O., de Ibarra, N.H., Philippides, A., and Collett, T.S. (2014). Head movements and the optic flow generated during the learning flights of bumblebees. J Exp Biol 217, 2633–2642. 10.1242/jeb.102897.

26. Schilstra, C., and van Hateren, J.H. (1998). Stabilizing gaze in flying blowflies. Nature 395, 654–654. 10.1038/27114.

27. Betts, C.R., and Wootton, R.J. (1988). Wing Shape and Flight Behaviour in Butterflies (Lepidoptera: Papilionoidea and Hesperioidea): A Preliminary Analysis. Journal of Experimental Biology 138, 271–288. 10.1242/jeb.138.1.271.

28. Jantzen, B., and Eisner, T. (2008). Hindwings are unnecessary for flight but essential for execution of normal evasive flight in Lepidoptera. Proceedings of the National Academy of Sciences 105, 16636–16640. 10.1073/PNAS.0807223105.

29. Pinkert, S., Friess, N., Zeuss, D., Gossner, M.M., Brandl, R., and Brunzel, S. (2020). Mobility costs and energy uptake mediate the effects of morphological traits on species’ distribution and abundance. Ecology 101, e03121. 10.1002/ecy.3121.

30. Suzuki, K., Nakamura, M., Kouji, M., and Yoshino, M. (2022). Revisiting the flight dynamics of take-off of a butterfly: experiments and CFD simulations for a cabbage white butterfly. Biol Open 11, bio059136. 10.1242/bio.059136.

31. Huang, H., and Sun, M. (2012). Forward flight of a model butterfly: Simulation by equations of motion coupled with the Navier-Stokes equations. Acta Mech Sin 28, 1590–1601. 10.1007/s10409-012-0209-1.

32. Sunada, S., Kawachi, K., Watanabe, I., and Azuma, A. (1993). Performance of a Butterfly in Take-Off Flight. J Exp Biol 183, 249–277. 10.1242/jeb.183.1.249.

33. Sikandar, U.B., Aiello, B.R., and Sponberg, S. (2025). Body oscillations couple with wing flapping to reduce aerodynamic power in wild silk moth flight. J. R. Soc. Interface. 22, 20250061. 10.1098/rsif.2025.0061.

34. Fei, Y.H.J., and Yang, J.T. (2016). Importance of body rotation during the flight of a butterfly. Physical Review E 93. 10.1103/PhysRevE.93.033124.

35. Sridhar, M., Kang, C.-K., and Landrum, D.B. (2019). Beneficial Effect of the Coupled Wing-Body Dynamics on Power Consumption in Butterflies. In AIAA Scitech 2019 Forum (American Institute of Aeronautics and Astronautics). 10.2514/6.2019-0566.

36. Johansson, L.C., and Henningsson, P. (2021). Butterflies fly using efficient propulsive clap mechanism owing to flexible wings. Journal of the Royal Society Interface 18. 10.1098/rsif.2020.0854.

37. Rutowski, R.L., and Warrant, E.J. (2002). Visual field structure in the Empress Leilia, Asterocampa leilia (Lepidoptera, Nymphalidae): dimensions and regional variation in acuity. J Comp Physiol A 188, 1–12. 10.1007/s00359-001-0273-7.

38. O’Carroll, D.C., and Wiederman, S.D. (2014). Contrast sensitivity and the detection of moving patterns and features. Philosophical Transactions of the Royal Society B: Biological Sciences 369, 20130043. 10.1098/rstb.2013.0043.

39. Nordström, K., Barnett, P.D., and O’Carroll, D.C. (2006). Insect detection of small targets moving in visual clutter. PLoS Biology 4, 0378–0386. 10.1371/journal.pbio.0040054.

40. Takeuchi, Y., Arikawa, K., and Kinoshita, M. (2006). Color discrimination at the spatial resolution limit in a swallowtail butterfly, Papilio xuthus. J Exp Biol 209, 2873–2879. 10.1242/jeb.02311.

41. Kraus, C.M., Jie, V.W., Hanslin, F.Ø., Grob, R., Beetz, M.J., Baird, E., and El Jundi, B. (2026). Stable neural coding of heading across locomotory modes by the insect compass system. Current Biology 36, 3405–3414.e5. 10.1016/j.cub.2026.05.063.

42. Cespedes, A., Penz, C.M., and DeVries, P.J. (2015). Cruising the rain forest floor: butterfly wing shape evolution and gliding in ground effect. Journal of Animal Ecology 84, 808–816. 10.1111/1365-2656.12325.

43. DeVries, P.J., Penz, C.M., and Hill, R.I. (2010). Vertical distribution, flight behaviour and evolution of wing morphology in Morpho butterflies. Journal of Animal Ecology 79, 1077–1085. 10.1111/j.1365-2656.2010.01710.x.

44. Le Roy, C., Amadori, D., Charberet, S., Windt, J., Muijres, F.T., Llaurens, V., and Debat, V. (2021). Adaptive evolution of flight in Morpho butterflies. Science 374, 1158–1162. 10.1126/science.abh2620.

45. Mathis, A., Mamidanna, P., Cury, K.M., Abe, T., Murthy, V.N., Mathis, M.W., and Bethge, M. (2018). DeepLabCut: markerless pose estimation of user-defined body parts with deep learning. Nature Neuroscience 2018 21:9 21, 1281–1289. 10.1038/s41593-018-0209-y.

46. Karashchuk, P., Rupp, K.L., Dickinson, E.S., Walling-Bell, S., Sanders, E., Azim, E., Brunton, B.W., and Tuthill, J.C. (2021). Anipose: A toolkit for robust markerless 3D pose estimation. Cell Reports 36, 109730. 10.1016/j.celrep.2021.109730.

47. Dang, A., Cerkvenik, U., Ilić, M., Pirih, P., Debevc, E., Briscoe, A.D., and Belušič, G. (2026). Graded opsin co-expression along the butterfly retina fine tunes the spectral sensitivity of a colour-opponent cell across the visual field. J Comp Physiol A 212, 245–258. 10.1007/s00359-025-01761-6.

48. Belušič, G., Ilić, M., Meglič, A., and Pirih, P. (2016). A fast multispectral light synthesiser based on LEDs and a diffraction grating. Scientific Reports 6, 1–9. 10.1038/srep32012.

49. Belušič, G., Ilić, M., Meglič, A., and Pirih, P. (2021). Red-green opponency in the long visual fibre photoreceptors of brushfoot butterflies (Nymphalidae). Proceedings of the Royal Society B 288. 10.1098/RSPB.2021.1560.

50. Peirce, J., Gray, J.R., Simpson, S., MacAskill, M., Höchenberger, R., Sogo, H., Kastman, E., and Lindeløv, J.K. (2019). PsychoPy2: Experiments in behavior made easy. Behav Res 51, 195–203. 10.3758/s13428-018-01193-y.

51. Juusola, M., Dau, A., Song, Z., Solanki, N., Rien, D., Jaciuch, D., Dongre, S.A., Blanchard, F., de Polavieja, G.G., Hardie, R.C., et al. (2017). Microsaccadic sampling of moving image information provides Drosophila hyperacute vision. eLife 6, e26117. 10.7554/eLife.26117.

52. Bomphrey, R.J., Walker, S.M., and Taylor, G.K. (2009). The Typical Flight Performance of Blowflies: Measuring the Normal Performance Envelope of Calliphora vicina Using a Novel Corner-Cube Arena. PLOS ONE 4, e7852. 10.1371/journal.pone.0007852.

